# Bifunctional enzyme SpoT is involved in biofilm formation of *Helicobacter pylori* with multidrug resistance by upregulating efflux pump *Hp1174* (*gluP*)

**DOI:** 10.1101/321026

**Authors:** Xiaoran Ge, Yuying Cai, Zhenghong Chen, Sizhe Gao, Xiwen Geng, Ya Li, Yan Li, Jihui Jia, Yundong Sun

**Author notes:** Xiaoran Ge and Yuying Cai contributed equally to this work. Address correspondence to Sun Yundong.

## Abstract

The drug resistance of *Helicobacter pylori (H. pylori)* is gradually becoming a serious problem. Biofilm formation is an important factor that leads to multidrug resistance in bacteria. The ability of *H. pylori* to form biofilms on the gastric mucosa has been known. However, there are few studies on the regulation mechanisms of *H. pylori* biofilm formation and multidrug resistance. Guanosine 3’-diphosphate 5’-triphosphate and guanosine 3’,5’-bispyrophosphate [(p)ppGpp] are global regulatory factors and are synthesized in *H. pylori* by the bifunctional enzyme SpoT. It has been reported that (p)ppGpp is involved in the biofilm formation and multidrug resistance of various bacteria. In this study, we found that SpoT also plays an important role in *H. pylori* biofilm formation and multidrug resistance. Therefore, it is necessary to carry out some further studies regarding its regulatory mechanism. Considering that efflux pumps are of great importance in the biofilm formation and multidrug resistance of bacteria, we tried to find if efflux pumps controlled by SpoT participate in these activities. Then, we found that *Hp1174* (glucose/galactose transporter, *gluP*), an efflux pump of the MFS family, is highly expressed in biofilm-forming and multi-drug resistance (MDR) *H. pylori* and is upregulated by SpoT. Through further research, we determined that *gluP* involved in *H. pylori* biofilm formation and multidrug resistance. Furthermore, the average expression level of *gluP* in clinical MDR strains was considerably higher than that in clinical drug-sensitive strains. Taken together, our results revealed a novel molecular mechanism of *H. pylori* tolerance to multidrug.

## INTRODUCTION

*Helicobacter pylori*, whose infection rate is over 50 percent throughout the world, is highly associated with a wide range of upper gastrointestinal diseases, especially gastric carcinoma(1). Contemporarily, the most common treatment method for this bacteria is called triple therapy, which consists of a proton-pump inhibitor and two antibiotics, namely, macrolides, nitroimidazoles or β-lactam(2). However, in recent years, eradication of *H. pylori* is becoming increasingly difficult because the rate of antibiotic resistance acquisition by *H. pylori* has generally increased(3). In addition, with the extended use of antibiotics, the appearance of multidrug-resistant (MDR) *H. pylori* strains, which are resistant to multiple antibiotics, has been reported(4).

There are numerous molecular mechanisms that contribute to multiple antibiotic resistance in the bacteria, including decreased drug permeability, efflux pumps, alteration or bypass of the drug target, production of antibiotic-modifying enzymes, and other physiological states, such as the formation of biofilms(5). Biofilms are communities of microorganisms anchored to a surface, which live in an extracellular matrix called extracellular polymeric substances (EPS) that they produce to form their immediate environment(6). The property that makes biofilms distinct from planktonic cells is their increased resistance to antimicrobial agents. Recently, it has become widely accepted that the biofilm plays an imperative part in the pathogenesis of some chronic human infections, as well as bacterial multidrug resistance(6, 7). Furthermore, it has already been reported that *H. pylori* has the ability to form biofilms in vitro(8). Then, in 2006, using scanning electron microscopy, Carron MA et al first proposed that *H. pylori* can form biofilms in vivo(9). The formation of biofilms in vivo is an important cause of *H. pylori* resistance to multiple antibiotics(10), while the stringent response to the stressful environment that lacks nutrients contributes to the resistance as well(11).

The stringent response is a bacterial stress response that controls bacterial adaptation to stress signals such as nutrient deprivation(12). In bacteria, the signal molecule (p)ppGpp, which is induced by diverse stresses, activates the stringent response(12, 13). The phenomenon of (p)ppGpp affecting bacterial multidrug resistance has already been reported for some other bacteria(14). Maisonneuve et al reported that under antibiotic stress, *E. coli* can produce rare cells that transiently become multidrug tolerant. In these rare cells, the level of (p)ppGpp was high(15). In addition, it has also been proven that (p)ppGpp can affect the formation of the bacterial biofilm. For example, Sugisaki et al have determined that the accumulation of the (p)ppGpp accelerated the formation of biofilms in *Bordetella pertussis* (16), and Li et al reported that the low level of (p)ppGpp contributed to the formation of biofilms in *Actinobacillus pleuropneumoniae S8*(17). Nevertheless, there are still no reports certifying the relationship between (p)ppGpp and the formation of biofilms in *H. pylori*.

In many bacteria, such as *E.coli*, (p)ppGpp is synthesized by two enzymes, RelA and SpoT(18), and SpoT is a bifunctional enzyme with both (p)ppGpp synthetase activity and hydrolase activity(18). However, there is only one member of RelA/SpoT family—SpoT—in the *H. pylori* genome(19); therefore, in *H. pylori*, we focused on whether SpoT is involved in *H. pylori* biofilm formation and multidrug resistance

There are a series of transport proteins in bacteria, which can acquire nutrients and extrude metabolic by-products, and some of these proteins are called efflux pumps(20). Efflux pumps have the ability to expel a broad range of antibiotics and have been recognized as significant components of multidrug resistance in many bacteria(21, 22), such as *H. pylori* (23–29). It has been demonstrated that efflux pumps work as one of the mechanisms that contribute to the antimicrobial resistance of biofilm(30), and evidence can be found in several microorganisms, such as *Pseudomonas aeruginosa* (31), *E. coli* (32) and *Candida albicans* (33). Moreover, Yonezawa et al reported that in *H. pylori* clinical strains, the expression of some efflux pump genes was elevated in biofilm cells compared to that in planktonic cells(34). While some efflux transporters have been detected in *H. pylori* 26695(19), their functions must be further studied, especially in *H. pylori* biofilm formation.

Considering that SpoT is a global regulator, we suppose that SpoT can influence the formation of biofilms and multidrug resistance by regulating the expression of the efflux pump in *H. pylori*.

## RESULTS

### SpoT is involved in *H. pylori* biofilm formation and multidrug resistance

As a global regulatory factor, SpoT has been proven to participate in the formation of bacterial biofilms(16, 17, 35), while no related studies have been done regarding *H. pylori*. Therefore, we analyzed the expression difference of *spoT* between biofilm-forming and planktonic cells by RT-PCR. The gene *spoT* is highly expressed in the latter (Fig. 1A). The (p)ppGpp assay shows that *H. pylori* in the biofilm produces more (p)ppGpp than the planktonic cells (Fig. 1B). By constructing a *spoT* mutant strains, we compared the biofilm of the wild-type and Δ*spoT* strains by GLSM (Fig. 1D) and SEM(Fig. 1E). The wild-type strains could form complete and compact biofilms on the NC membrane, whereas for the biofilm of the Δ*spoT* strains, the bacteria were not packed tightly enough and the biofilm matrix was not complete, and some cavities could be seen (Fig. 1E). LIVE/DEAD cell viability assays show that Δ*spoT* strains form a lighter biofilm than the wild-type strains do (Fig. 1C/D).

**Fig. 1:**
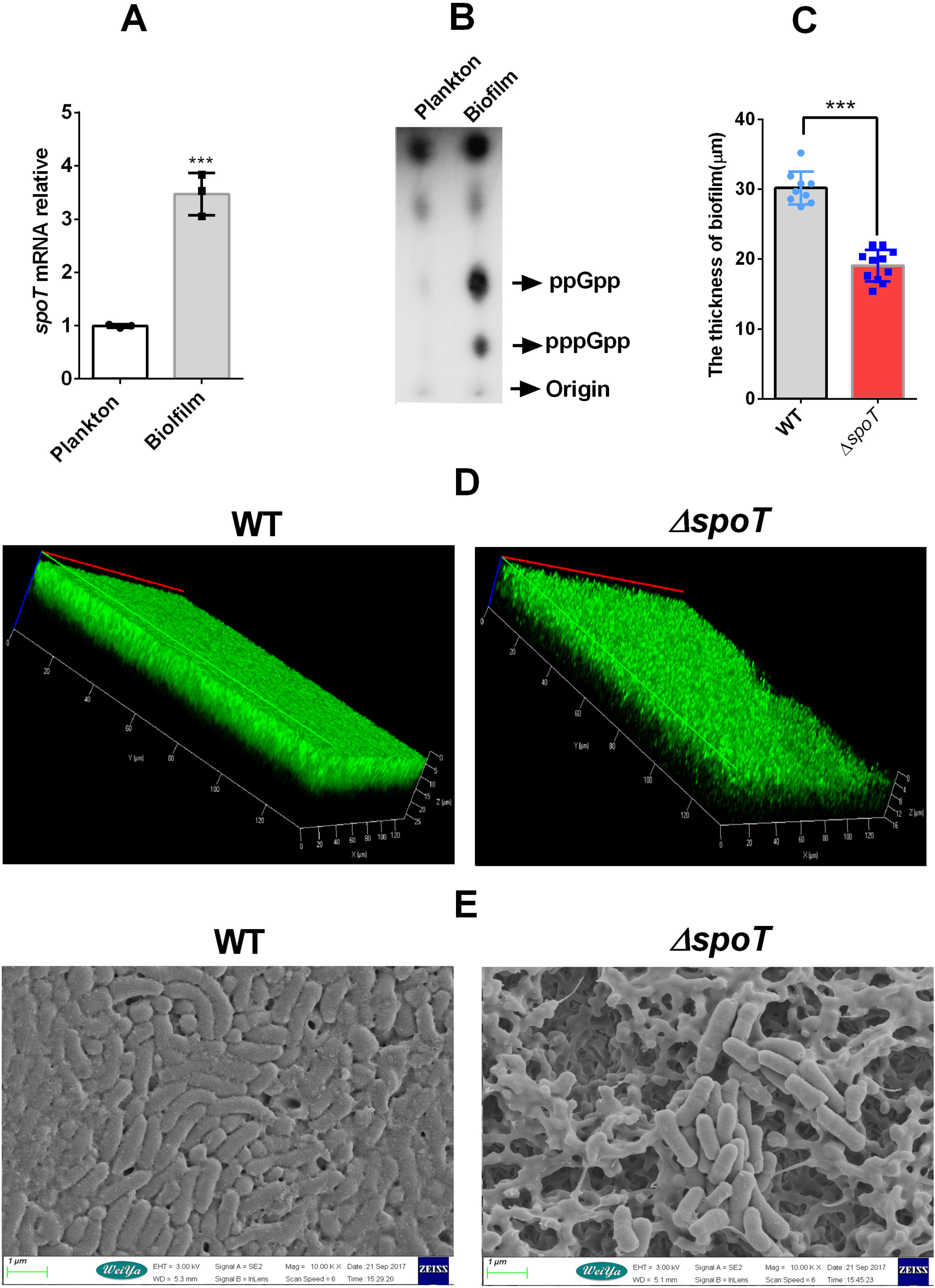
SpoT is involved in *H. pylori* biofilm formation. A: qRT-PCR mRNA expression of SpoT in biofilm-forming and planktonic cells. The signals were normalized to the 16S rRNA levels. Data are the means ±SEM from three independent experiments. Significance by paired Student’s *t* test: ***, *P* <0.001. B: (p)ppGpp was induced in the biofilm-forming cells, but not in the planktonic cells.^32^P-labeled nucleotides of the formic acid extracts of *H. pylori* were detected by thin-layer chromatography. Planktonic *H. pylori* were grown to the exponential phase. C: Comparing the biofilm thickness between WT and Δ*gluP* strains, data from (D), The data presented are the means ±SEM from three independent experiments. Significance was determined by paired Student’s *t* test:***, *P* <0.001. D: Confocal laser scanning microscopy (CLSM) images of the WT and *ΔspoT* strains biofilms. Cells stained with membrane-permeable Syto 9 (green) and membrane-impermeable PI (red) were visualized by confocal microscopy. E: Scanning electron microscopy (SEM) images of WT and Δ*spoT* strains biofilms. The biofilm used in this experiment is a mature biofilm grown on a nitrocellulose membrane for three days, and planktonic bacteria were from early exponential phase (OD600, approximately 0.4–0.5).

According to the MIC of the wild-type strains and the Δ*spoT* strains, for planktonic cells, the Δ*spoT* strains are apparently more sensitive to various antibiotics than the wild type, and ciprofloxacin is not included (Tab. 2). With regard to the biofilm-forming cells, after knocking out *spoT*, the cell resistance to various antibiotics is reduced, especially for penicillin G (Tab. 2). Then, we treated the wild-type and Δ*spoT* strains with the MIC of the antibiotics (CLA, AMO, TET and MET). The growth inhibitory curve demonstrates that the growth ability of the Δ*spoT* strains are obviously inhibited (Fig. 2).

**Fig. 2.**
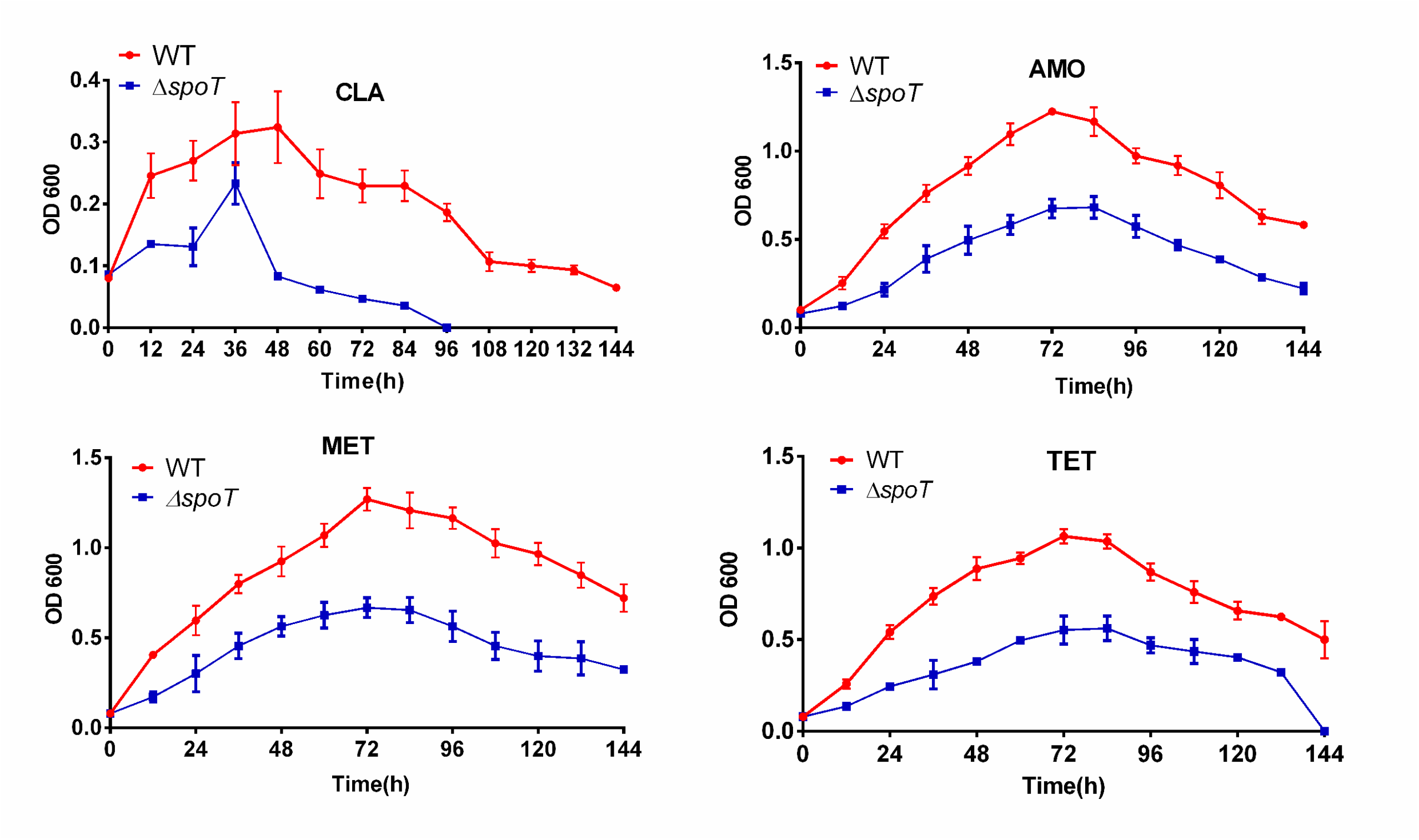
Comparison of growth inhibition curves characteristics of WT and Δ*spoT* strains with various antibiotics. (AMO (0.125 μg/ml), CLA (0.125 μg/ml), MET (0.5 μg/ml), TET (0.25 μg/ml)). Data are the means ±SEM from three independent experiments. Significance was determined using paired Student’s t test: ***, *P* <0.001.

### Comparing the efflux capacity of wild and Δ*spoT* strains

As SpoT is involved in the formation of the biofilm and multidrug resistance of *H. pylori*, and an efflux pump contributes to them as well(22, 30), we inferred that SpoT could possibly carry out those functions by regulating the expression of efflux pump. Therefore, we compared the efflux activity of the wild-type and Δ*spoT* strains. The results reveal that whether in biofilm-forming or planktonic cells, the inactivation of SpoT caused a distinct increase in the accumulation of Hoechst 33342, notably for planktonic cells, and the fluorescence values of the Δ*spoT* strains were >3-fold greater than those of the WT. These results demonstrate that the efflux activity of Δ*spoT* strains to Hoechst 33342 is weaker than that of the wild-type strains (Fig. 3), which suggests that SpoT is likely to regulate the efflux pump gene expression of *H. pylori*.

**Fig. 3.**
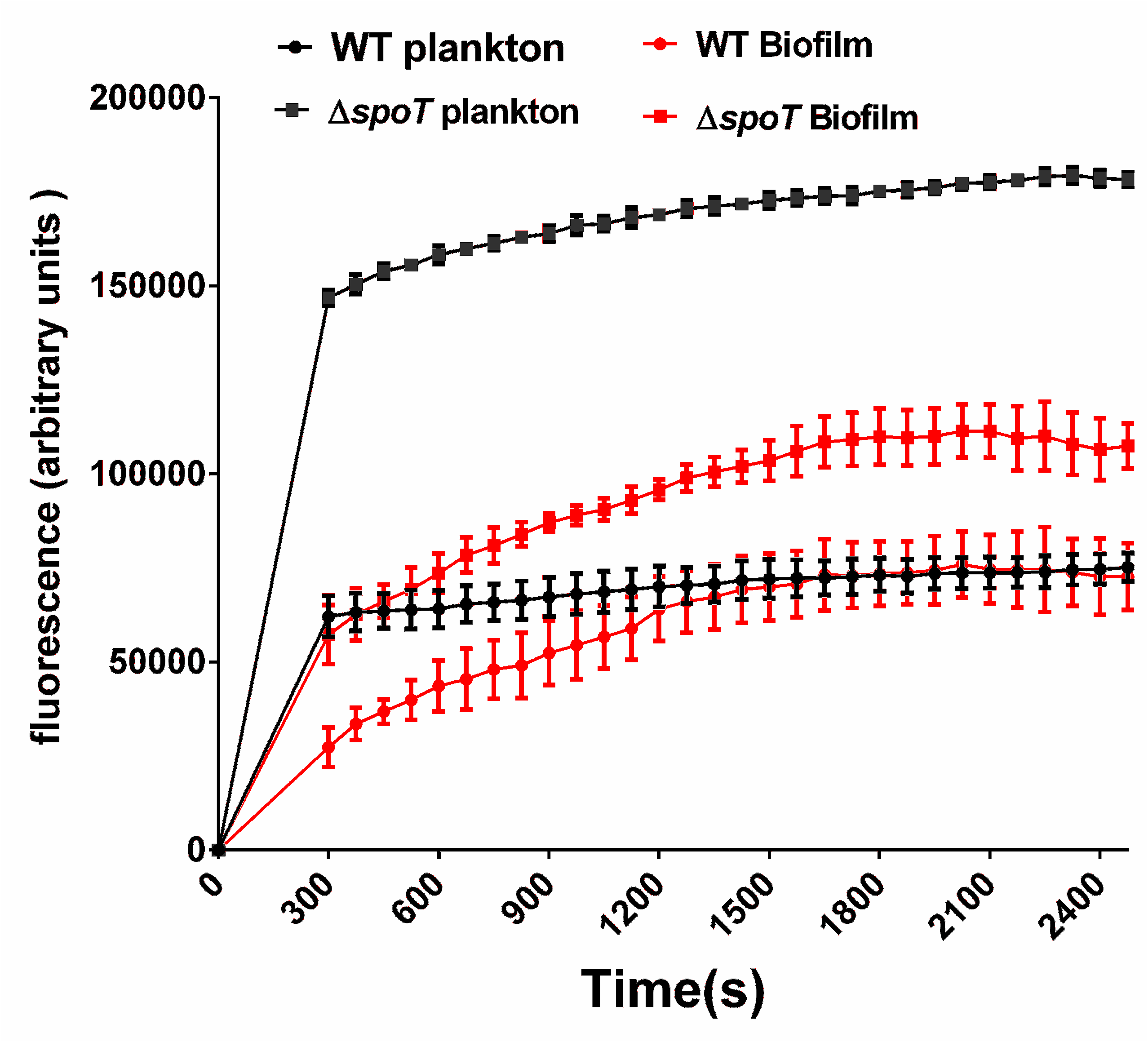
Comparison of the accumulation of H33342 (2.5 M) in biofilm and planktonic cells of WT and Δ*spoT* strains. The fluorescence intensity was recorded at the excitation and emission wavelengths of 350 and 460 nm, respectively, over a 30-min incubation period. The data presented are the means ±SEM from three separate experiments. A paired Student’s *t* test was performed to compare the accumulation of Hoechst 33342 between the WT and Δ*spoT* strains. ***, *P* <0.001.

### Screening efflux pumps involved in the formation of biofilms and multidrug resistant in *H. pylori* by qRT-PCR

There are six families of bacterial efflux pumps: (1) the ATP-binding cassette (ABC) superfamily; (2) the major facilitator superfamily (MFS); (3) the multidrug and toxic compound extrusion (MATE); (4) the small multidrug resistance (SMR) family; (5) the resistance-nodulation-division (RND) superfamily; and (6) the drug metabolite transporter (DMT) superfamily(30). It is generally agreed that ABC, MFS and RND family play important roles in gram-negative bacteria(30). As extensive studies regarding RND family in *H. pylori* have been conducted so far(8, 25, 29, 34), we analyzed the expression difference of some efflux pumps belonging to the MFS and ABC family between biofilm-forming and planktonic cells by using qRT-PCR (Tab. 3); As seen from the table, the expression levels of two particular genes had greater than three-fold increases within biofilm-forming cells compared those in to planktonic cells: *hp1181* and *hp1174* (*gluP*). Furthermore, we analyzed the mRNA expression difference of these two genes in multidrug-resistant *H.pylori* (MDR-H, artificially selected), and only *gluP* was highly expressed, compared to that in the sensitive strains (Fig. 4). Considering that SpoT shows high expression in MDR-H (Fig. 4) strains and biofilm-forming cells, we proposed that SpoT may participate in the formation of biofilm and multidrug resistance of *H. pylori* by upregulated *gluP*.

**Fig. 4.**
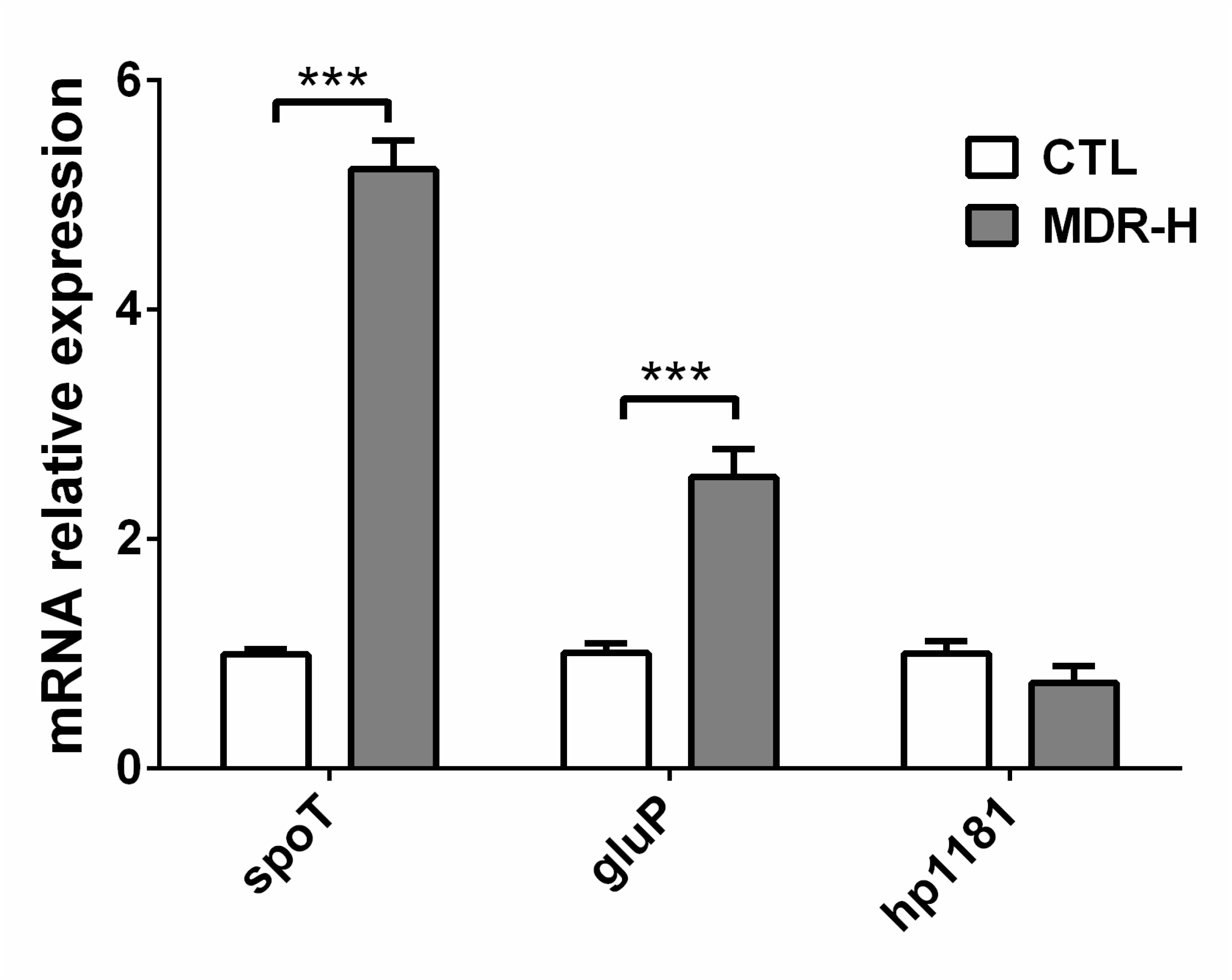
qRT-PCR analysis of the mRNA levels of *spoT*, *gluP* (*hp1174*) and *hp1181* in MDR-H strains (selected artificially) compared to those of WT. The signals were normalized to the 16S rRNA levels. Data are the means±SEM from three independent experiments. Significance was determined by paired Student’s *t* test: **, *P* <0.01; ***, *P* <0.001.

### SpoT regulates GluP expression

To verify if GluP is regulated by SpoT, we chose three antibiotics (CLA, AMO and MET) to treat the wild-type strains and Δ*spoT* strains. First, we used three kinds of antibiotics with different concentrations to stimulate the wild-type and Δ*spoT* strains for 10 min; then we treated the wild-type strains and the Δ*spoT* strains at a specific concentration of the three antibiotics along a time gradient. The results of qRT-PCR showed that AMO and MET can both induce the expression of *gluP* in wild-type strains compared to that in the control group (wild-type strains without antibiotics treatment), doing so in a concentration-and time-dependent manner, while in the Δ*spoT* strains, GluP can barely be induced by these two antibiotics (Fig. 5). However, CLA can hardly induce the expression of *gluP*, and the sensitivity of wild and Δ*spoT* strains to CLA is similar according to MIC data. This suggests that *gluP* may not be involved in the resistance of *H. pylori* to CLA. In short, the results above indicate that SpoT upregulates GluP to cope with specific antibiotics stress.

**Fig. 5.**
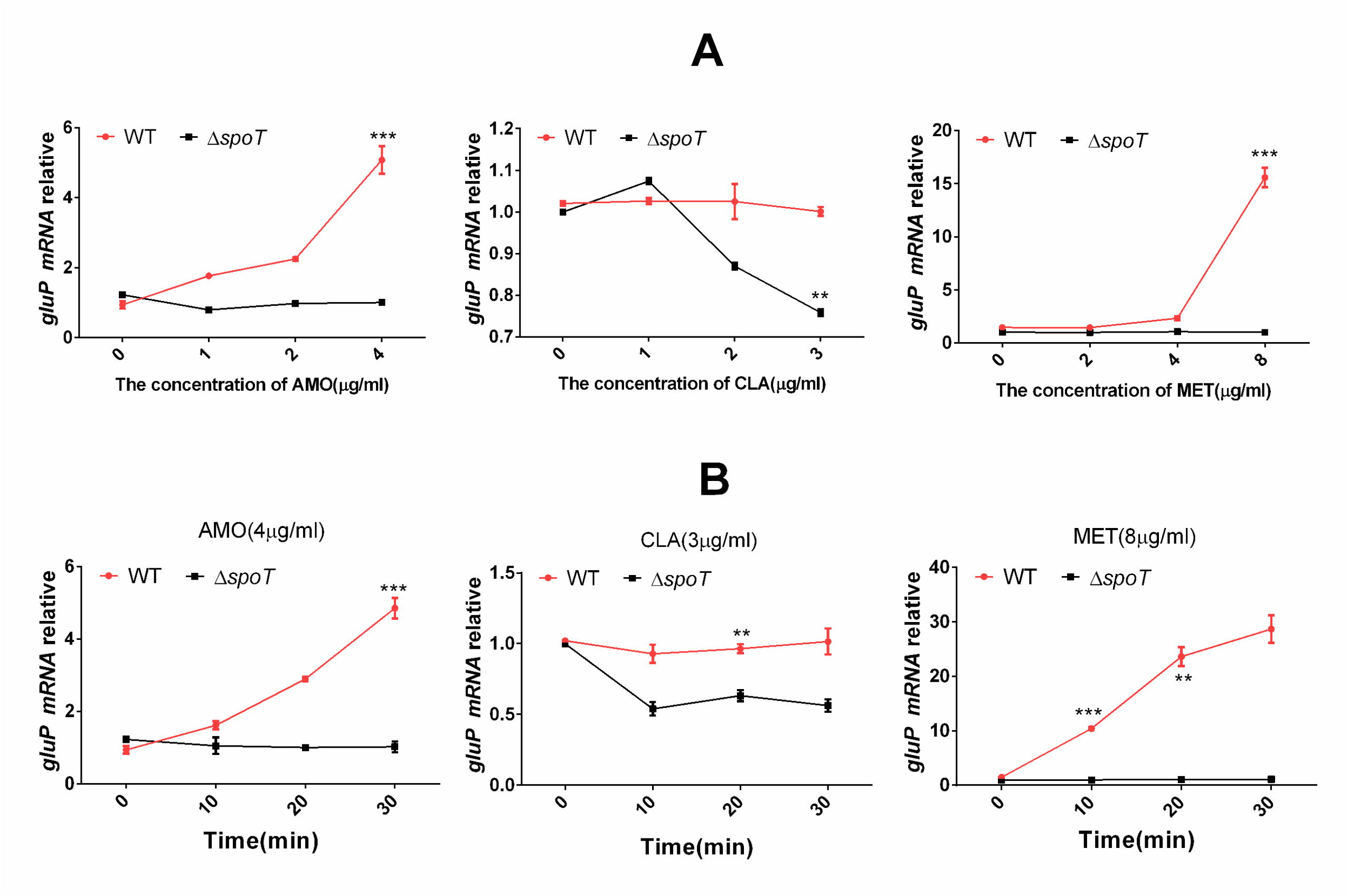
qRT-PCR analysis of the mRNA levels of *gluP* in the WT and Δ*spoT* strains. WT and Δ*spoT* strains exposed to different drug concentrations for 10 min (A) and exposed to special drug concentrations for different time periods (B). The results were compared to the WT without drug treatment (CTL). The signal was normalized to the 16S rRNA levels. Data are the means ±SEM from three independent experiments. Significance was determined by paired *t* test: **, *P* < 0.01; ***, *P* <0.001.

### GluP involved in *H. pylori* efflux

GluP, a glucose transporter, is responsible for glucose transport in *H. pylori* (19). In addition, the analysis of structure demonstrates that GluP is an efflux pump belonging to MFS family, which suggests that GluP is likely to take part in drug efflux in *H. pylori*. Therefore, we successfully constructed *gluP* mutant strains and compared the efflux capacity of wild-type strains and Δ*gluP* strains. The results reveal that whether in biofilm-forming and planktonic cells, the inactivation of GluP caused a distinct increase in the accumulation of Hoechst 33342, especially for planktonic cells, and the fluorescence values of the Δ*gluP* strains were >3-fold greater than those of the wild-type strains. These results showed that Δ*gluP* strains had lower efflux capacity for Hoechst 33342 than that of wild-type strains, both in planktonic and biofilm-forming cells (Fig. 6).

**Fig. 6.**
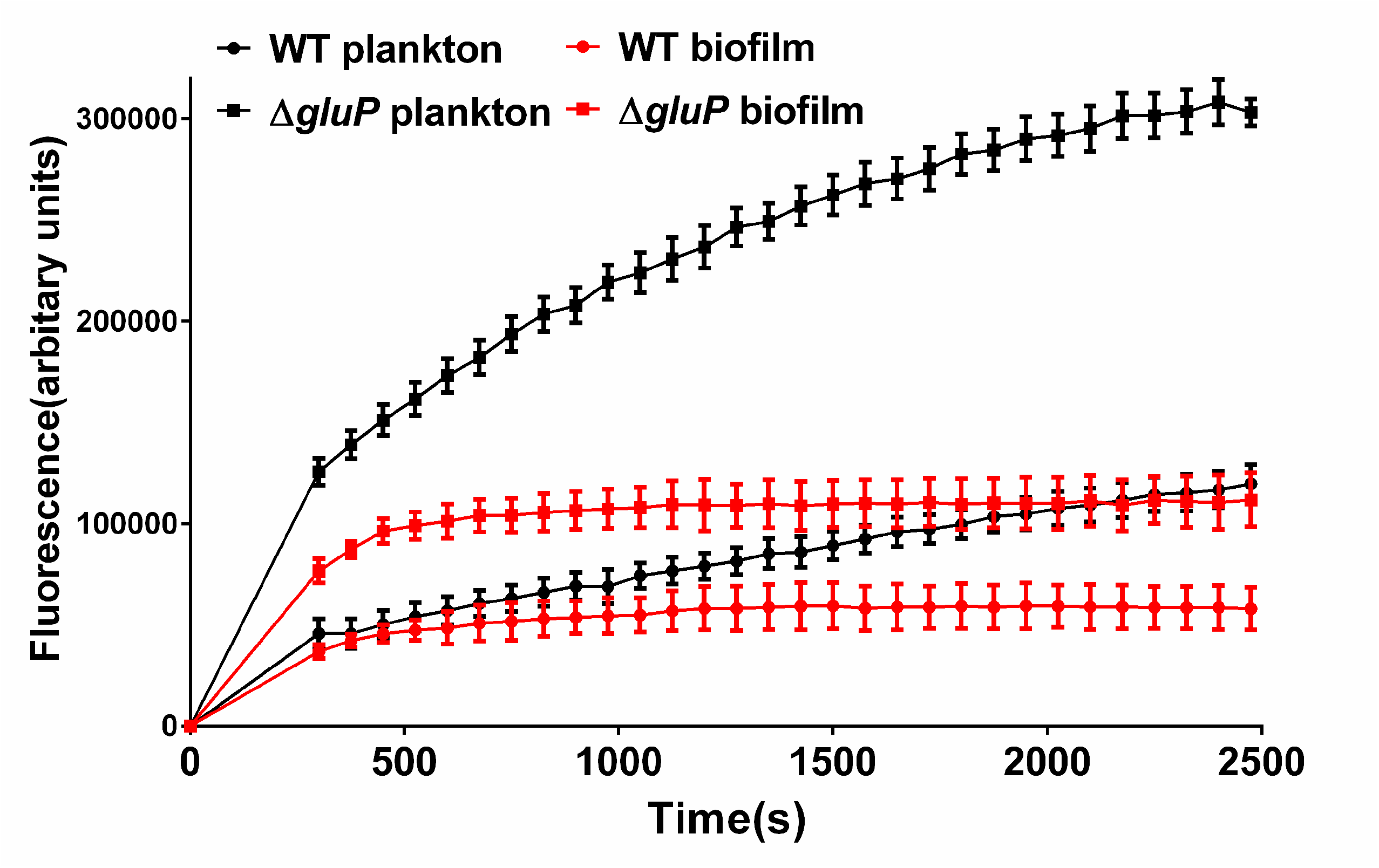
Comparison of the accumulation of H33342 (2.5 M) in biofilm and planktonic cells of WT and Δ*gluP* strains. The fluorescence intensity was recorded at the excitation and emission wavelengths of 350 and 460 nm, respectively, over a 30-min incubation period. The data presented are the means ±SEM from three separate experiments. Paired Student’s *t* test was performed to compare the accumulation of Hoechst 33342 between the WT and Δ*gluP* strains. ***, *P* <0.001.

### Glup is involved in *H. pylori* biofilm formation and multidrug resistance

Studies have reported that efflux pumps participate in bacterial biofilm formation(30), so we compared biofilms of wild-type strains and Δ*gluP* strains by SEM. The images showed that compared with the wild-type strains, bacteria in the Δ*gluP* strain biofilm were not tightly packed, and the biofilm matrix was incomplete and showed more cavities (Fig. 7A). By GLSM, LIVE / DEAD cell viability assays showed that the Δ*gluP* strains form a thinner biofilm (Fig. 7B/C). According to the MIC of the wild-type strains and Δ*gluP* strains from planktonic cells (Tab. 2), the Δ*gluP s*trains are apparently more sensitive to various antibiotics than the wild-type strains, except for clarithromycin and ciprofloxacin. After knocking out *gluP* in MDR-H strains, its resistance to drugs also dropped significantly, especially for amoxicillin (Tab. 2). In addition, for those biofilm-forming cells, after knocking out *gluP*, its sensitivity to various antibiotics increased (Tab. 2).

**Fig. 7.**
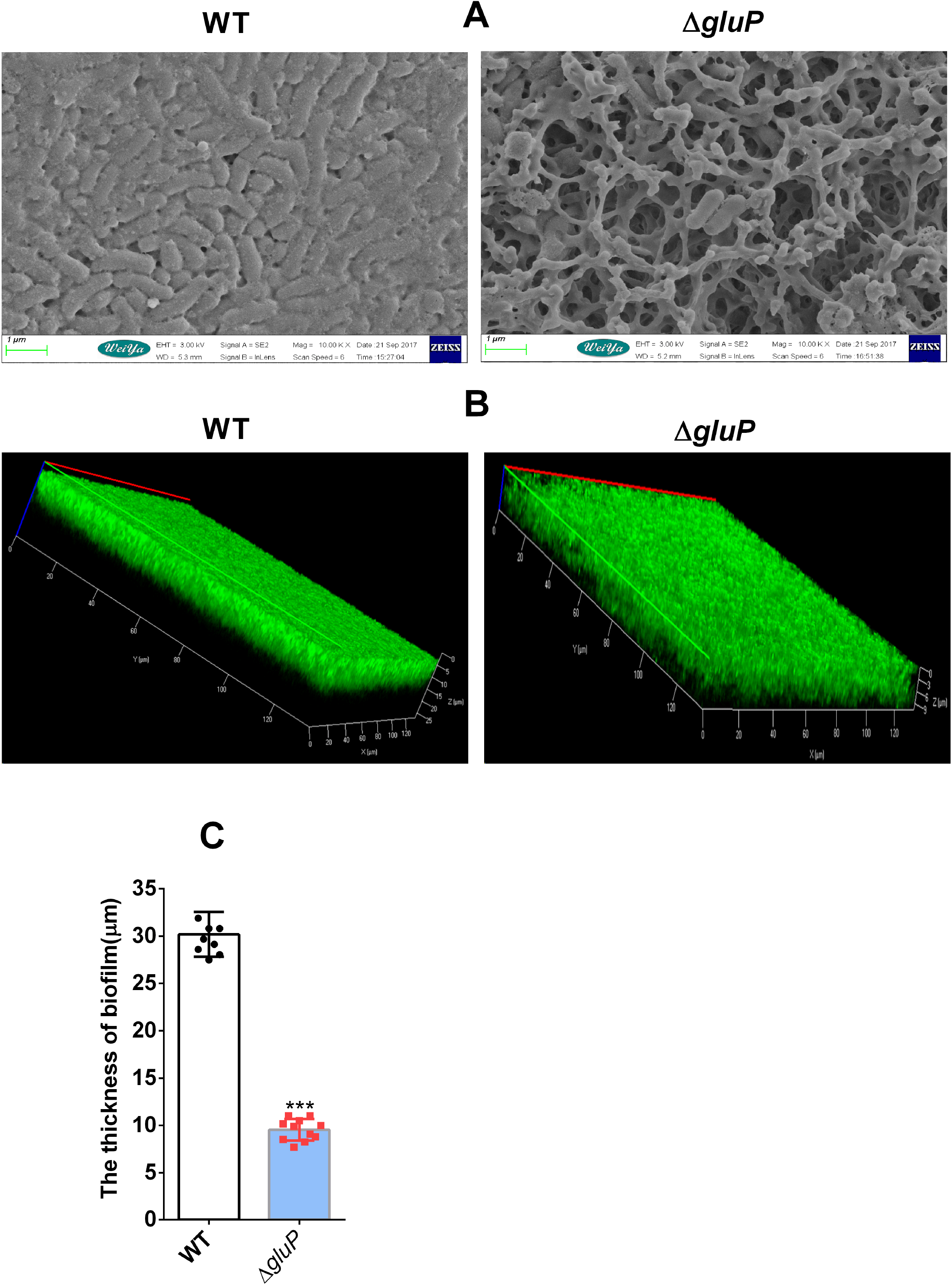
GluP is involved in *H. pylori* biofilm formation. A: Scanning electron microscopy(SEM) images of WT and Δ*gluP* strain biofilms. B: Confocal laser scanning microscopy (CLSM) images of the WT and Δ*gluP* strains biofilms. Cells stained with membrane-permeable Syto 9 (green) and membrane-impermeable PI (red) were visualized by confocal microscopy. C: Comparing the biofilm thickness between WT and Δ*gluP* strains, data from (B), The data presented are the means ±SEM from three independent experiments. Significance was determined by paired Student’s *t* test:***, *P* <0.001. The biofilm used in this experiment is a mature biofilm grown on a nitrocellulose membrane for three days, while the planktonic *H. pylori* was from the early exponential phase (OD600, approximately 0.4–0.5).

We treated wild-type strains and Δ*gluP* strains separately with the MIC (of wild strains) of three antibiotics (CLA, AMO and MET) and then observed their growth inhibition curve. The results showed that compared with the wild-type strains, the growth ability of the Δ*gluP* strains was significantly inhibited by AMO and MET, but not CLA (Fig. 8). The above results suggest that GluP may not be involved in planktonic *H. pylori’s* tolerance to clarithromycin, which is consistent with the results of MIC.

**Fig. 8.**
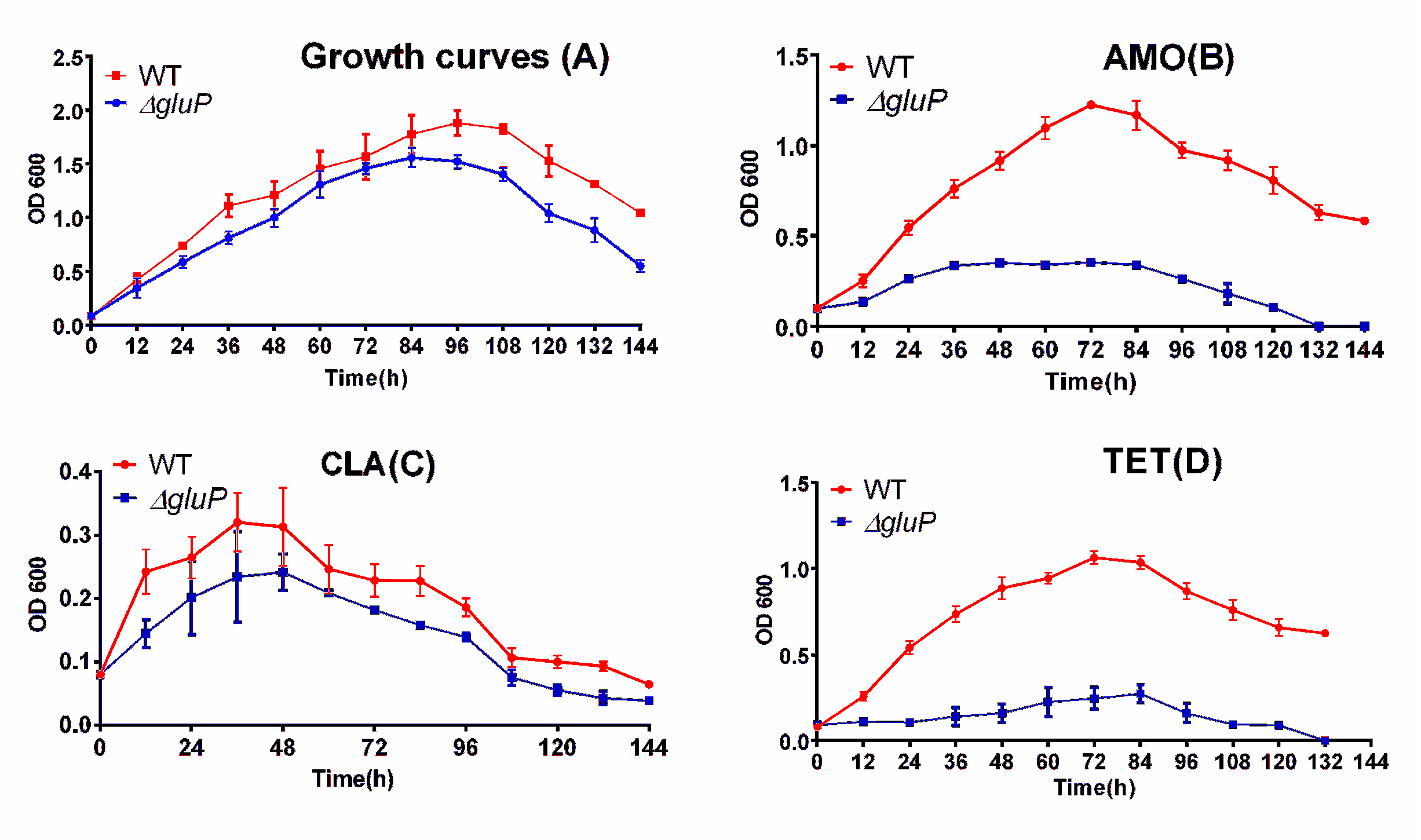
Comparison of growth curves and growth inhibition curves characteristics of WT and Δ*gluP* strains. A: Growth curves of the WT and Δ*gluP* strains. B, C and D: Growth inhibition curves of the WT and Δ*gluP* strains exposed to AMO (0.125 μg/ml), CLA: (0.125 μg/ml) and MET (0.5 μg/ml). Data are the means ±SEM from three independent experiments. Significance was determined by paired Student’s *t* test: ***, *P* <0.001.

The relative mRNA expression levels of *gluP* were assessed by qRT-PCR in clinical MDR strains and clinical sensitive strains. The average expression levels of *gluP* in clinical MDR strains were significantly higher than those in sensitive ones (Fig. 9).

**Fig. 9.**
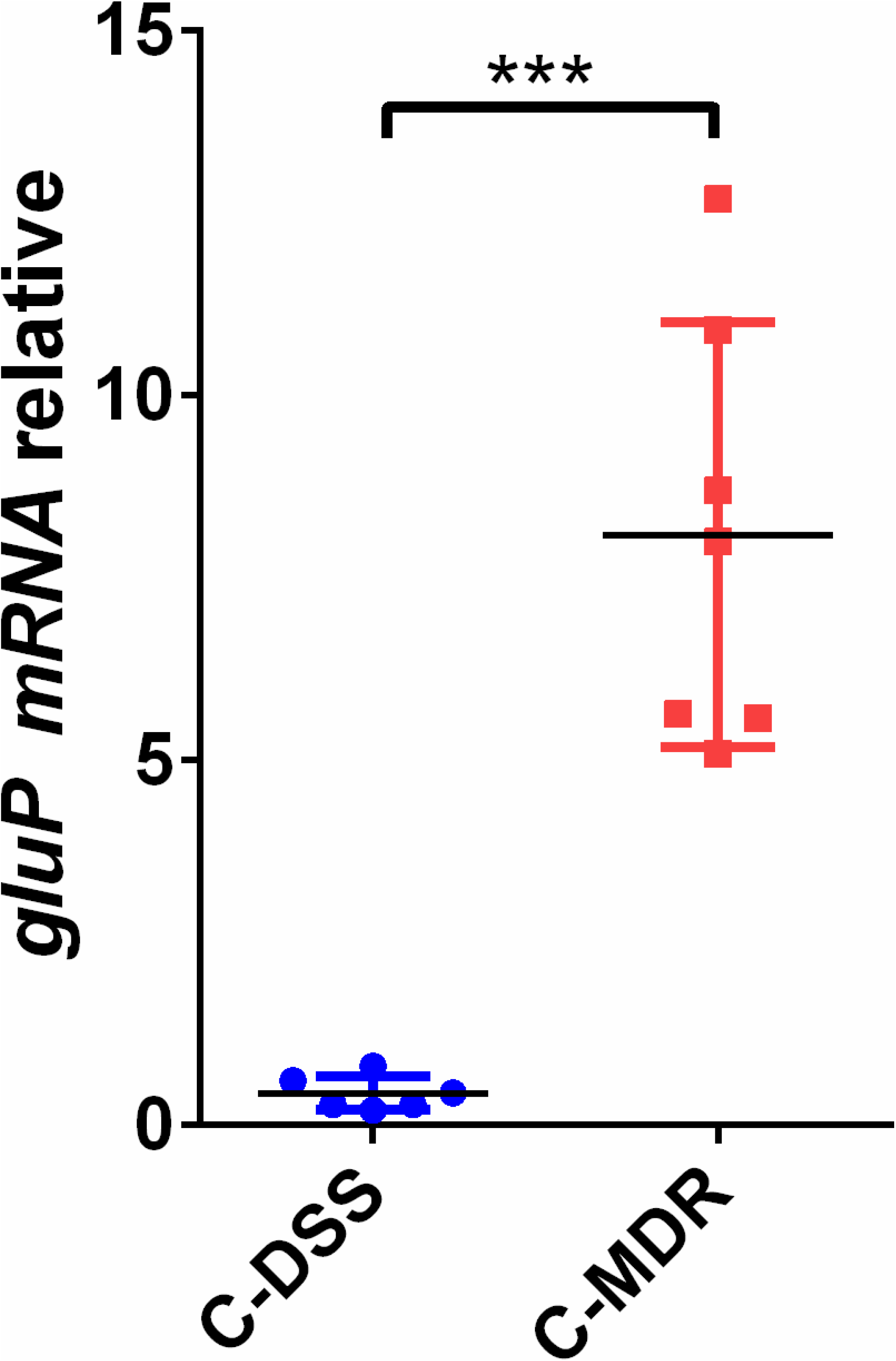
Analysis of *gluP* expression differences in clinical drug-sensitive strains (C-DSS) and clinical multidrug-resistant strains (C-MDR) by qRT-PCR. The results were compared to the value of *gluP* in the WT. The signal was normalized to the 16S rRNA levels. Data are the means ±SEM from three independent experiments. Significance was determined by paired Student’s *t* test: ***, *P* <0.001.

## DISCUSSION

Recently, antibiotic resistance acquired by *H. pylori* has generally increased, and the formation of biofilms in vivo is an important cause leading to its multidrug resistance(10). In this research, we found that SpoT is involved in the formation of biofilm in *H. pylori*. While it has been reported that efflux pump takes part in the formation of biofilms and multidrug resistance(30), we found that *gluP* is involved in the biofilm formation and multidrug resistance of *H. pylori*. Further analysis showed that the expression of *gluP* was upregulated by SpoT. Considering that SpoT is a global regulator, our study partly explained the molecular mechanism of SpoT involved in the biofilm formation of *H. pylori* and multidrug resistance.

Antibiotics, as signaling molecules(36), can induce bacteria to produce (p)ppGpp and lead to a stringent response(14). Studies have shown that (p)ppGpp is involved in bacterial tolerance to various antibiotics such as penicillin(37), vancomycin(38) and ampicillin(39), (p)ppGpp is synthesized in *H. pylori* from SpoT(19). Our previous studies discovered that (p)ppGpp modulates the expression of *H. pylori* efflux transporters(40), accompanied with the importance of efflux pump to multidrug resistance(21); thus, we wondered if (p)ppGpp participates in multidrug resistance of *H. pylori*. This research confirmed that inference.

The formation of biofilm is another important factor that causes multidrug resistance in bacteria(7). Biofilms are communities of microorganisms anchored to a surface, living in an extracellular matrix called extracellular polymeric substances (EPS), which are produced by the organisms to form their immediate environment(7). Bacteria with biofilms have strong resistance to antibiotics and host immune defenses(6). It has been reported that (p)ppGpp is involved in the formation of biofilm in various bacteria, but the function of (p)ppGpp during this process varies from species to species. Several previous studies showed that the lack of (p)ppGpp resulted in decreased biofilm formation in bacteria, such as *Enterococcus faecalis* (35), *Vibrio cholerae* (41) and *Bordetella pertussis* (16). However, other studies proved that (p)ppGpp-synthase deletion mutants of *Actinobacillus pleuropneumoniae* (17) and *Francisella novicida* (42) could produce significantly more biofilms than the wild type. Since SpoT is the only gene that plays a part in the synthesis of (p)ppGpp in *H. pylori*, its ability to synthesize (p)ppGpp is totally lost after the knockout of SpoT, consequently lowering its ability to adapt to stressful environments. In this study, we induced *H. pylori* to form a biofilm through nutrient starvation, because nutrition deficiency is a significant factor that induces bacteria to form biofilms. After *spoT* knockout, the ability of *H. pylori* to form biofilms was greatly reduced.

The formation of biofilms is an important cause of multidrug resistance in bacteria, and biofilms lead to continuous chronic infections, which add to the difficulty in providing a clinical cure(6). Drug resistance mechanisms include the following: poor diffusion of antibiotics through the biofilm polysaccharide matrix, physiological changes due to the slow growth rate and starvation responses that form persistent cells, phenotypic changes of the cells forming the biofilm, the expression of efflux pumps, and so on(7). Drug efflux is a key mechanism of drug resistance in gram-negative bacteria(21). Microorganisms regulate the internal environment to adapt to their outer circumstances by excluding poisonous substances (microbial agents, metabolite and quorum sensing chemical signals) via efflux pumps(22). Normally, the expression of efflux pumps in biofilm-forming bacteria is higher than that in planktonic cells(30). For example, in *P. aeruginosa*, the mechanism for tolerance to collistin is the upregulation of the MexAB-OprM efflux pump in biofilms(43). Moreover, in *E. coli*, when growing in biofilms and exposed to several antibiotics, AcrAB-TolC efflux pump-encoded genes are upregulated(44). In addition, the expression levels of the acrA and acrB genes are observed to increase in Salmonella biofilm cells(45). It has been reported that the RND family of efflux pumps are highly expressed in *H. pylori* biofilms(34), and recent studies have shown that *hp1165* and *hefA* are highly expressed in clinically isolated *H. pylori* biofilms(46). In our research, the expression of several efflux pumps is high in *H. pylori* biofilms. Some further studies regarding *gluP* found that it took part in the formation of *H. pylori* biofilms. After *gluP* knockout, the *H. pylori* biofilm matrix is damaged and unable to form a well-structured biofilm. Some studies confirmed that the extremely low biofilm formation in the wild type can be observed in *E. coli* mutants without emrD, emrE, emrK, acrD, acrE and mdtE efflux pump-encoding genes(47). Efflux pumps are involved in biofilm formation, which is possibly due to the export or import of some substances related to forming biofilms. *gluP* is a glucose/galactose transporter, belonging to the major facilitator superfamily (MFS)(19), which is mainly responsible for the physiological uptake of sugars, such as D-glucose, into *H. pylori*. D-Glucose is involved in the synthesis of bacterial exopolysaccharides, while polysaccharides account for a major fraction of the biofilm matrix(48). Therefore, the deletion of *gluP* may affect the synthesis of the *H. pylori* biofilm matrix. We found that the matrix of the *H. pylori* biofilm was incomplete after gluP knockout, even if the concentration of glucose in the medium was increased (supplementary, Fig S1). In addition, the biofilm matrix can limit the transport of some antimicrobial agents to cells within the biofilm(7).

For the time being, extensive studies concerning the regulatory mechanism of efflux pumps have been conducted, such as two-component signal (TCS) transduction systems, local repressors and global response regulators(49). In *E. coli*, (p)ppGpp can also regulate the expression of the efflux pump of YojI(50). There is one possible mechanism that explains how (p)ppGpp indirectly mediates global changes in the transcriptional level, such as by reducing the RpoD competitiveness for the core RNA polymerase. (p)ppGpp releases RpoD from the RNA polymerase and shifts the use of the alterative sigma factors(51). It is known that the *H. pylori* genome only includes two alterative sigma factors (σ^54^ and σ^28^)(19). According to the research of Niehus and coworkers, the promoter sequence of σ^54^-dependent genes is TTTGCTT(52). By analyzing the upstream sequence of the putative ATG start codon of the ORFs of *gluP*, we discovered a possible conserved sequence (TTTGCAT) that was recognized by σ^54^ (supplementary, Fig. S2), which suggests that *gluP* could be regulated by σ^54^. In further studies, antibiotic treatment could not induce high *gluP* expression in the σ^*54*^ mutant strains (supplementary, Fig. S3). Therefore, SpoT may upregulate the expression of *gluP* by controlling σ^54^, but detailed mechanisms require more studies.

In conclusion, this research has discovered a new mechanism regarding the formation of the biofilms and multidrug resistance in *H. pylori*. It must be admitted that further analysis is required to prove specific mechanisms of how SpoT regulates *H. pylori* biofilm formation. On account of the present data, our research provides evidence and clues for the clinical treatment of drug-resistant strains and epidemiological investigation.

## MATERIALS & METHODS

### Bacterial strains, media, growth conditions, clinical isolation of *H. pylori*

*H. pylori* 26695, which was kindly provided by Dr. Zhang Jianzhong (Chinese Disease Control and Prevention Center), was used in this study. The strain was maintained at –80°C in Brucella broth (Difco, Detroit, MI) with 20% (vol/vol) glycerol and 10% fetal bovine serum (FBS). The strain was cultured under a microaerobic environment (5% O_2_, 10% CO_2_, and 85% N_2_) at 37°C on Brucella agar plates containing 7% lysed sheep blood. The liquid culture medium for *H. pylori* consisted of Brucella broth (BB) containing 10% fetal bovine serum (FBS), and the cells were incubated in a shaker set at 120 rpm at 37°C. The mutant strains were cultured on agar plates with kanamycin (Sigma-Aldrich, St. Louis, MO) at 30 μg/ml. The *E. coli* strain used was TOP10 (TransGen Biotech, Beijing, China), and grew in Luria–Bertani medium at 37°C.

Eight multidrug-resistant and eight sensitive clinical isolates were obtained from patients, including those with gastritis, gastric ulcers, duodenal ulcers and gastric cancer, at Qiannan People’s Hospital (Guizhou Province). All the patients provided informed consent before examination. MDR *H. pylori* from the patients could not be killed with repeated standard triple therapy treatment. Isolated *H. pylori* strains were identified using universally accepted phenotypic tests: typical morphology on Gram-stained smears and positive urease, oxidase and catalase tests.

### Assessment of susceptibility to antibiotics

The minimal inhibitory concentrations (MICs) of all the clinical and standard strains for various antibiotics were determined by the Etest and the agar dilution method reported by Osato et al.(53). The bacteria (optical density at 600 nm [OD600], 0.8) were inoculated on an agar plate containing 2-fold dilutions of antibiotics. All the plates were incubated at 37°C under microaerobic conditions, and MIC values were determined. The determination of biofilm-forming bacterial MICs was similar to that for the planktonic bacteria. The biofilm, which was attached to the NC membrane, was incubated in a liquid medium containing different concentrations of various antibiotics for 12 h. After being washed three times with PBS, the biofilm bacteria were suspended in liquid culture medium, and this liquid was inoculated on an agar plate to determine the MIC values.

### Construction of biofilm

The biofilm could be cultivated in two kinds of conditions: the static condition or the continuous flow condition. There were many ways to cultivate biofilms under the static condition, among which we used the “colony biofilm” (described in previous article(54, 55)) with slight modifications. To allow the adherence of *H. pylori* at the interface, 20 μl of bacteria were inoculated at 5×10^7^ cells ml^-1^ onto a nitrocellulose membrane (approximately 1×1 cm^2^), which was placed on an agar plate. The agar plate was cultured in a microaerobic environment (5% O_2_, 10% CO_2_, and 85% N_2_) at 37°C for 3 days.

### Scanning electron microscopy

To observe the *H. pylori* biofilm, SEM (scanning electron microscopy) was used. The samples for SEM were prepared along the following standard procedures. Planktonic bacteria were fixed with glutaraldehyde after centrifugation. For the biofilm-forming bacteria, the samples were gently washed with autoclaved PBS three times to remove planktonic bacteria. The biofilms were fixed in 2.5% glutaraldehyde at 4°C for 2 h and then washed three times with cacodylate buffer and dehydrated through a series of graded ethanol solutions (25%, 50%, 75%, 95% and 100%). Subsequently, the samples were freeze-dried, sputter coated with gold, and observed by a scanning electron microscope.

### Confocal laser scanning microscopy (CLSM)

To determine bacterial shape and viability, planktonic *H. pylori* were stained with membrane-permeable and membrane-impermeable fluorescent dyes from the Live/Dead BacLight Bacterial Viability kits (Molecular Probes, Invitrogen, USA). Then, they were observed by confocal microscopy, which was performed according to a previous study(40). For biofilm-forming cells, the NC membrane with the biofilm was removed from the agar plate and then gently washed three times with PBS. Subsequently, the NC membrane was stained in a 12-well plate with 1 ml Syto 9 dye for 20 mins in the dark conditions and gently washed three times with PBS. After that, the NC membrane was fixed on a slide, covered with a coverslip and subsequently analyzed with a confocal laser scanning microscope (Leica TCS SP5; Leica Microsystems GmbH, Wetzlar, Germany). Syto 9 is a green fluorescent membrane-permeable dye that labels all bacteria by the staining nucleic acid, whereas PI is a red fluorescent membrane-impermeable dye that only labels bacteria with damaged membranes.

### Detection of (p)ppGpp accumulation patterns

The (p)ppGpp production was assayed according to the previous literature with slight modification(40). The treatment of the planktonic bacteria was the same as previously described. For the biofilm bacteria, first, they were washed with PBS, then diluted to an OD600 of 0.2, and incubated for an additional 2 h. When all the strains reached an OD600 of approximately 0.3, the samples of each culture plate were centrifuged at 10,000 rpm for 5 min, and resuspended in 250 μl of liquid culture medium.^32^P (Amersham) was added to 100 μCi ml^-1^, and the cultures were labeled for 2 h at 37°C. Then, 50 μl of each sample was added to an equal volume of 2 M formic acid. Afterward, at least four freeze-thaw cycles were conducted. The acid extracts were briefly centrifuged, and the supernatant fluids were spotted onto the polyethyleneimine-cellulose plates (Sigma-Aldrich), dried, and developed in 1.5 M KH_2_PO_4_ (pH 3.4) for approximately 2.5 h. The results were obtained under phosphor screen scanning (Bio-Rad).

### Extraction of RNA and Real-time Quantitative PCR (qRT-PCR)

The total RNA was extracted using TRIzol reagent (Invitrogen, Carlsbad, CA). To reverse transcribe the total RNA, the PrimeScript RT Reagent kit with gDNA Eraser (TaKaRa) was used. Primers are shown in Table 1. In the 20 μl qPCR mixtures, 5 μl of the resulting cDNA, which was already diluted, 10 μl of SYBR Premix Ex TaqTM (TaKaRa, Otsu, Shiga, Japan), 0.8 μl of the primer mixture, and 4.2 μl of double-distilled water were added. Then, RT-PCR was done using the ABI Prism 7000 Sequence detection system (Applied Biosystems, Carlsbad, CA) for 1 cycle at 95°C for 30 s and 40 cycles at 95°C for 5 s and 60°C for 31 s. Dissociation curve analysis was performed to verify the product homogeneity. The 16S rRNA amplicon was used as an internal control for data normalization. Changes in the transcript level were determined by applying the relative quantitative method (ΔΔC_T_). The threshold cycle (C_T_) values from all three biological replicates for each strain were compiled.

### Construction of *hp1174* (*gluP*) mutant strains

The plasmids pILL570 and pUC18K2 were kindly provided by Agnès Labigne (Unité de Pathogénie Bactérienne des Muqueuses, Institut Pasteur). The construction of the *gluP* mutant strains was identical to the construction of the Δ*spoT* strains, as described in the previous literature (56). Briefly, the genome of *H. pylori* 26695 was extracted to serve as a template. Primers are listed in Table 1. The gene *hp1174* was destroyed with the insertion of the nonpolar aphA-3 gene encoding a kanamycin resistance cassette.

**TABLE 1.**
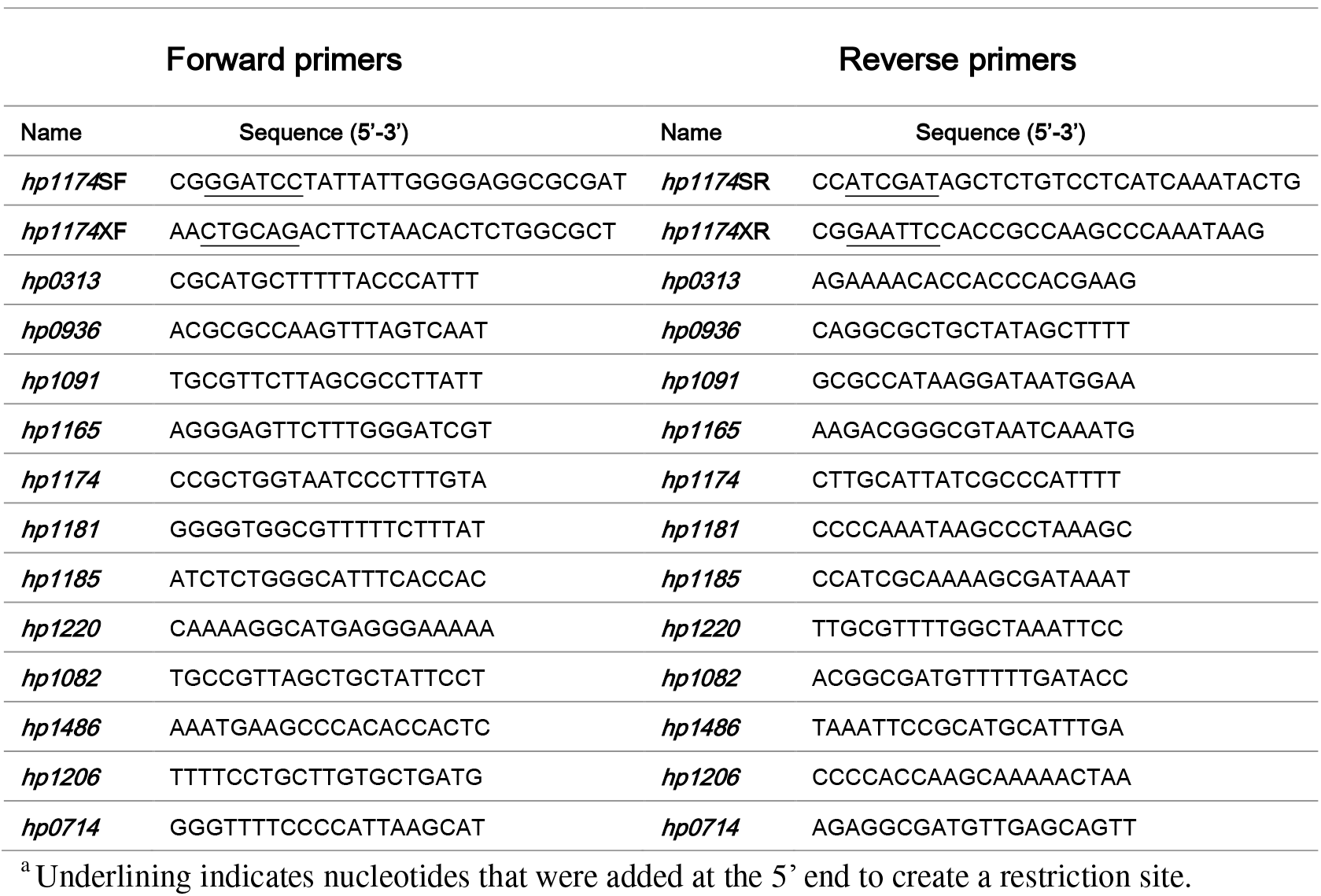
Primers used in this study^a^

**TABLE 2.**
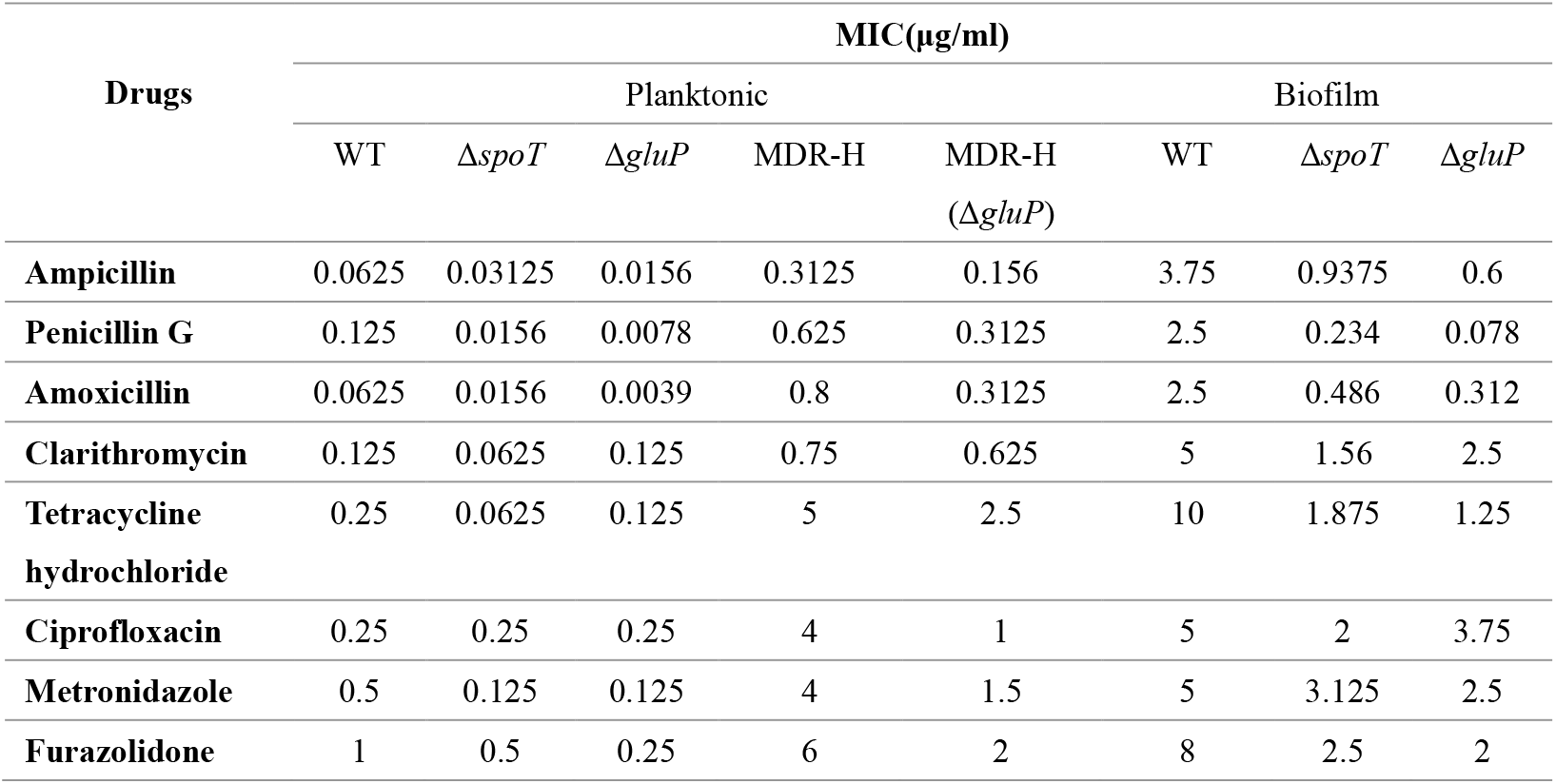
MICs determined for WT, Multidrug-resistant漏MDR漐strains, SpoT and GluP mutant strains in biofilm-forming and planktonic *H.pylori*

**Table 3.** **qRT-PCR** analysis of efflux pumps expression difference between the biofilm-forming and the planktonic *H.Pylori*

| Efflux pump families. | Locus Tag | Predicted Function | Fold Value Biofilm/Plankter | P value Significance |
| --- | --- | --- | --- | --- |
| **The Major Facilitator Superfamily (MFS)** | HP0313 | Nitrite | 0.46±0.14 | * |
|  | HP0936 | Proline/betaine | 1.08±0.17 |  |
|  | HP1091 | Alpha-ketoglutarate | 0.58±0,03 | ** |
|  | HP1165 | Multi-drug efflux | 2.66±0.86 |  |
|  | HP1174 | Glucose/galactose transporter | 5.75±0.68 | *** |
|  | HP1181 | multidrug efflux | 3.34±0.31 | *** |
|  | HP1185 | Sugar efflux | 0.45±0.27 | * |
| **The ATP-binding Cassette (ABC) Superfamily** | HP1220 | Multi-drug efflux | 2.99±0.32 | ** |
|  | HP1082 | Multi-drug efflux | 2.02±0.25 |  |
|  | HP1486 | Multi-drug efflux | 1.11±0.29 |  |
|  | HP1206 | Multi-drug efflux | 1.93±0.17 | ** |

### Determination of growth curves and growth inhibition curves

The determination of the growth curves was consulted in the previous literature(40). The growth profiles were monitored in Brucella broth with a preliminary OD600 of 0.08 and then cultured for another 144 h at 37°C with shaking. Records were taken every 12 h by determining the OD600 of the test strains. The values stated are the mode values from at least three biological replicates performed in at least three independent occasions.

To analyze the growth inhibition curve, the *H. pylori* strains were inoculated into Brucella broth with a preliminary OD600 of 0.08, which also contains different kinds of antibiotics (MIC of wild strains), and then cultured for another 144 h at 37°C with shaking. Each experiment was repeated at least three times.

### H33342 accumulation assay

For the planktonic bacteria, the accumulation assay method was performed as described previously(40). The biofilm bacteria was first rinsed off with PBS and subsequently suspended in PBS, with the final OD600 of the suspensions being adjusted to 0.1. Then, 180 μl of this liquid and 20 μl of H33342 (25 μM) was added to each well in a 96-well plate. Recording was started 5 min after the addition of H33342. The excitation and emission were measured at 355 nm and 460 nm, respectively, using FLUO star Optima (Aylesbury, UK). Readings were taken every 75 s for 30 cycles, and the raw data were analyzed by Excel. Each experiment was repeated at least three times.

### Generation of multidrug-resistant *H. pylori* (MDR-H)

To induce the chloramphenicol resistant strains, *H. pylori* 26695 was cultivated on the agar plate which contained 0.5× minimum inhibitory concentration (MIC) chloramphenicol for 48–72 h under micro-aerobic environments at 37°C, as described in the previous article(57). The resistant colonies were incubated with repeated doubling of the chloramphenicol concentration until no colony was seen. Colonies were maintained on the agar plates containing 4×MIC of tetracycline, ampicillin, penicillin G, and erythromycin. Then, the colonies were incubated for 48–72 h under a microaerobic environment.

### Statistical analysis

Data are presented as the means ± standard errors of the means (SEM). Statistical significance was determined using an unpaired Student’s *t* test, and the *P* values were corrected by the Sidak-Bonferroni method for multiple comparisons. *P* values of <0.05 were considered statistically significant. The results were analyzed using GraphPad Prism software (GraphPad Software Inc., La Jolla, CA, USA).

## CONFLICTS OF INTEREST

All authors have no conflict of interest.

## ACKNOWLEDGMENTS

The present research was supported by the National Natural Science Foundation of China (No. 81471991, 81671978, 81460314, 81374101 and 81571960).

We declare that we have no conflicts of interest.

